# Dynamic Entrainment: A deep learning and data-driven process approach for synchronization in the Hodgkin-Huxley model

**DOI:** 10.1101/2023.04.17.537224

**Authors:** Soheil Saghafi, Pejman Sanaei

## Abstract

Resonance and synchronized rhythm are important phenomena and can be either constructive or destructive in dynamical systems in the nature, specifically in biology. There are many examples showing that the human’s body organs must maintain their rhythm in order to function properly. For instance, in the brain, synchronized or desynchronized electrical activities can lead to neurodegenerative disorders such as Huntington’s disease. In this paper, we adopt a well known conductance based neuronal model known as Hodgkin-Huxley model describing the propagation of action potentials in neurons. Armed with the “data-driven” process alongside the outputs of the Hodgkin-Huxley model, we introduce a novel ***Dynamic Entrainment*** technique, which is able to maintain the system to be in its entrainment regime dynamically by applying deep learning approaches.

## 1 Introduction

Resonance or an increase in amplitude is a phenomenon that occurs when the frequency of an applied periodic force and the natural frequency of the system which the force acts on, are the same or close to each other. In that case, the system oscillates with a greater amplitude, which can either be destructive or constructive [1–6]. For instance, a footbridge with an equal resonant frequency to that of human walking might collapse. In 1850, a powerful thunderstorm caused the oscillation of Angers Bridge, while a group of marching soldiers tried to cross by swaying to maintain their balance. This event created resonance and most likely caused the bridge to collapse with the soldiers [7]. The instability of the London Millennium Bridge or so called “Wobbly Bridge”, which was induced by pedestrians walking on it during the opening day, enforced the closure of the bridge for almost two years for further repairs and modifications to make the bridge more stable [8].

Synchronization is a general term that refers to a process of interaction between two or more systems which are able to adjust their pace over time. Depending on the situation, synchronization or desynchronization of two or more dynamical systems can be beneficial. The history of synchronization goes back to Christiaan Huygens, a well known Dutch mathematician, physicist, and engineer, who invented the first functional pendulum clock in 1656 to help sailors to navigate their location on the globe. During setting up the clocks, Huygens discovered a pair of pendulum clocks hanged by a wood beam, which initially swung entirely out of phase over a period of time, eventually synchronized and remained in a lockstep phase [9, 10]. After Huygens, the study of synchronization events has been attracting researchers from a wide range of fields, including mechanical and electrical engineering as well as mathematics, physics, and biology [11–13]. In biology, one might refer to fireflies in South-East Asia, which synchronize their flashes, even though each of them prefers to flash at a different frequency. Flashing makes the fireflies coupled to each other strongly enough such that hundreds or even thousands of fireflies can flash simultaneously in a split second [14].

There are extensive examples in biology exhibiting that the human’s body organs must maintain their rhythm in order to function properly. For instance, palpitation or arrhythmia occurs when an electrical signal fires from a wrong place at a wrong time which in turn causes the heart to beat out of rhythm [15, 16]. In the brain, synchronized or desynchronized electrical activities can lead to neurodegenerative disorders like Huntington’s disease [17, 18]. Suprachiasmatic Nucleus, often known as the body clock, is a tiny region in the brain that controls the 24-hour circadian rhythm of human’s body. An improper entrainment to the light or dark cycle through Suprachiasmatic Nucleus can result in delayed or advanced sleep phase disorders [19–24].

Entrainment, which is a type of synchronization, refers to a phase-locked situation of an oscillation to an external periodic input [25–34]. N:M patterns are used to describe the type of entrainment defined as the number of input oscillations (N) that are phase-locked to the number of output oscillations (M). The number of input and output cycles are equal in 1:1 entrainments [35]. In this paper, we introduce a technique to find the ideal time interval for preserving the 1:1 entrainment regime while utilizing the force system. We adopt a well known conductance based neuronal model known as Hodgkin-Huxley model describing the propagation of action potentials in neurons [36]. Armed with the “data-driven” process alongside the outputs of the Hodgkin-Huxley model, we introduce a novel ***Dynamic Entrainment*** technique, which is able to maintain the system in its entrainment regime dynamically by applying Artificial Intelligence (AI) methods. Although, some AI techniques like Reinforcement Learning [37–40] do not depend on data and learn from the environment, other AI techniques like any machine learning or deep learning models utilize a synthetic dataset in their training process [41–44]. Dynamic Entrainment is categorised as the second group as it requires data in its procedure and the “data-driven” process [45, 46] assists to extract information from the data directly, which can be either the output of the mechanistic model or experimental results. The drawback of employing experimental data is that there is not sufficient amount of data compared to those generated by the mechanistic model. However, as soon as the data-driven model is developed based on the synthetic data originated from the mechanistic model, one must apply it on the experimental data, to infer more realistic results.

This paper is organized as follows: in section 2, we introduce the mechanistic model (a conductance-based model that describes the electrical activity of neurons in the brain) and the structure of the deep learning model, which are used throughout the paper. In section 3, we explain the data-driven process and define the concept of input-predictors along with output-responses which are the key components in the deep learning training process. Afterwards, we introduce the notion of Dynamic Entrainment and we present how well this technique is able to keep the pattern in a 1:1 entrainment regime by adapting the pace of the applied force to the system. Finally, we conclude and summarize our modeling results in Section 4 and provide some insight into real-world applications as well as potential future improvements.

## 2 Model description

### 2.1 Mechanistic model

The mechanistic model we consider in this paper is a conductance-based model describing the electrical activity of neurons introduced by Hodgkin and Huxley in 1952. The model explains the ionic mechanisms underlying the initiation and propagation of action potentials in the squid giant axon [36]. Hogkin-Huxley type models use an electric equivalent circuit to describe the excitability of the cell membrane. Figure 1*A* represents the equivalent circuit corresponding to the Hodgkin-Huxley equations [47]. The figure shows the cell membrane capacitance *C*, ionic conductances *g*_*y*_, where *y ∈ {Na, K, L}* for sodium *Na*, potassium *K* and leak *L*, the equilibrium potential or reversal potential *E*_*y*_, and the applied current *I*_*app*_(*t*) as a function of time *t*.

**Fig. 1:**
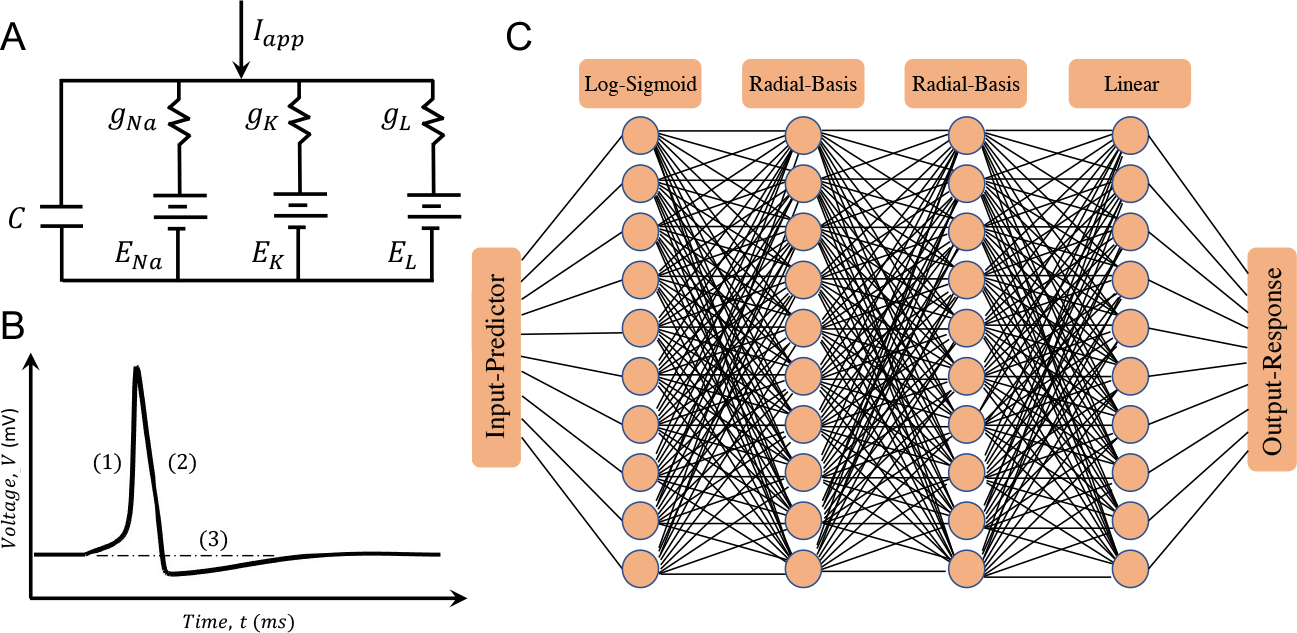
(*A*) Equivalent circuit underlying the Hodgkin-Huxley equations [47] showing the cell membrane capacitance *C*, ionic conductances *g*_*y*_, where *y ∈ {Na, K, L}* for sodium *Na*, potassium *K* and leak *L*, the equilibrium potential or reversal potential *E*_*y*_, and the applied current *I*_*app*_(*t*) as a function of time *t*. (*B*) Schematic of the cell membrane potential *V* (mV) versus time *t* (ms) with an action potential and three main phases: (1) depolarization of the cell membrane through activation of the voltage-dependent sodium channels, (2) repolarization of the cell membrane through inactivation of the sodium channels and activation of potassium channels, and (3) refractory period time during which the cell membrane is not able to fire or generate another action potential. (*C*) The neural network structure used in this paper. It consists of four hidden layers, each includes ten neurons. The first hidden layer is a Log-Sigmoid activation function, followed by two Radial Basis and a Linear activation functions.

At steady state, the inside of the cell is more negatively charged compared to the outside, leading to a hyperpolarized membrane potential *V* of around *−*60 or *−*70 mV. The cell membrane acts as a capacitor *C* that separates charges, however there are channels in the membrane that conduct certain ions. For sodium ions, the extracellular concentration is higher than that of intracellular, while for potassium ions the opposite is true. Due to these ionic concentration gradients, when sodium channels are open *Na*^+^ ions tend to flow into the cell and depolarize the membrane potential, whereas *K*^+^ ions flow out of the cell and re-polarize the membrane potential when potassium channels are open. Figure 1*B* represents the schematic of the cell membrane potential *V* (mV) versus time *t* (ms) with an action potential and three main phases: (1) depolarization of the cell membrane through activation of the voltage-dependent sodium channels, (2) repolarization of the cell membrane through inactivation of the sodium channels and activation of potassium channels, and (3) refractory period during which time the cell membrane is not able to fire or generate another action potential.

The mentioned ionic currents are described by Ohm’s Law, for example, for sodium ions we have *I*_*Na*_ = *g*_*Na*_*m*^3^*h*(*V − E*_*Na*_), where *I*_*Na*_ is the sodium current, *g*_*Na*_ is the maximal conductance of the sodium channel and *E*_*Na*_ is the equilibrium or reversal potential at which no current flows. The gating variable *m* describes whether the sodium channels are closed (*m →* 0) or open (*m →* 1), and the other gating variable *h* models whether the sodium channels are inactivated (*h →* 0) or de-inactivated (*h →* 1). Similar equations can explain the ionic current for potassium and leak ions *I*_*K*_ and *I*_*L*_, respectively (see (1)). Note that, the potassium channel has a single gating variable *n*, since it does not inactivate. In addition, the leak channel *L* is passive and thus there is not any gating variable associated with it.

The equations representing the Hodgkin-Huxley model are

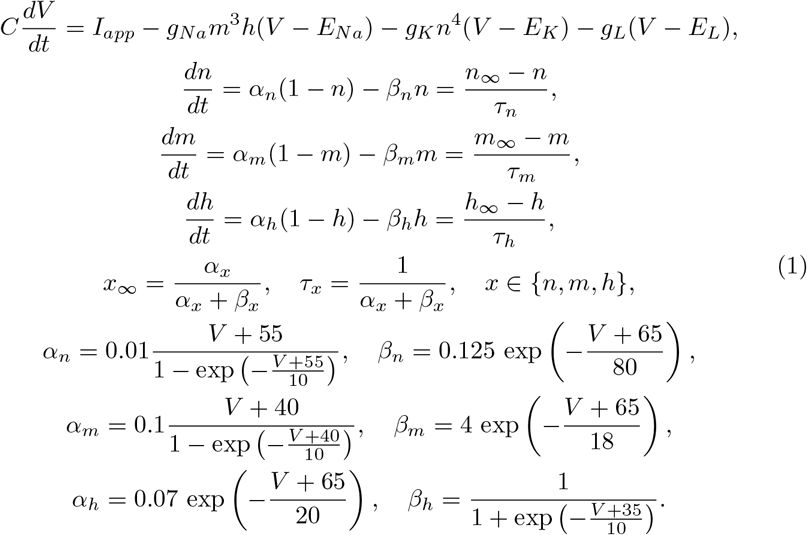

Throughout the paper, the parameter values and gating functions are *C* = 1 *µ*F/cm^2^, *g*_*Na*_ = 120 mS/cm^2^, *g*_*K*_ = 36 mS/cm^2^, *g*_*L*_ = 0.3 mS/cm^2^, *E*_*Na*_ = 55 mV, *E*_*K*_ = *−*77 mV and *E*_*L*_ = *−*54.4 mV. The applied current value depends on the time interval of the injected current, i.e. *I*_*app*_ = *Amp µ*A/cm^2^ if 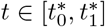 with length 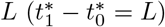 and *I*_*app*_ = 0 elsewhere. Note that 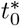 and 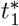 are the initiation and termination of the applied force, respectively.

The current-balance equation (1) originates from Kirchhoffâs Current Law (the algebraic sum of all currents entering and exiting a node in a circuit must equal zero) and says the sum of the capacitive and ionic currents must be equal to the applied current *I*_*app*_. The gating variables *n, m* and *h* in (1) are based on a two-state channel model with *x* being the fraction of open channels and 1 *− x* the fraction of closed channels, with voltage-dependent rates of closed channels opening *α*_*x*_ and open channels closing *β*_*x*_, where *x ∈ {n, m, h}*. Note that the gating variable equations can be rewritten in terms of the steady-state activation *x*_*∞*_ and time constant *τ*_*x*_ functions as shown in (1). By solving this system of ODEs in (1), we obtain the membrane potential over time *V* (*t*), as shown in Fig. 1*B*. The dynamics of the voltage-dependent gating variables, with *m* being on a faster time scale than *n* and *h*, create positive and negative feedbacks in the cell membrane. This results in a stable periodic solution that corresponds to the repetitive firing of action potentials.

### 2.2 Predictor model

The data-driven technique is one of the decision making procedure that relies on analyzing the provided data that is usually obtained directly from the experiment. Due to lack of sufficient experimental data, we use the mechanistic model in (1), to generate the data. This data can have different types: discrete or continues; supervised or unsupervised. Depending on the type, one can choose either any machine learning or deep learning approaches to analyze the data. Although, a very deep neural network model has a higher accuracy compared to other machine learning models, in order to have a trade off between the accuracy and the speed of the training process, here we construct a shallow neural network model. As shown in Fig. 1*C*, this model has four hidden layers including ten neurons each, a Log-Sigmoid, two Radial Basis and a linear activation functions.

## 3 Results

In this paper, we introduce Dynamic Entrainment which is a technique using data-driven process alongside the outputs of the mechanistic model for a dynamical system to train the deep learning approach in order to build a predictor to determine the best time of activeness of the force to keep the system in its 1:1 entrainment regime. This section consists of two subsections: Section 3.1 focuses on the data-driven process and generating a comprehensive synthetic dataset for building the predictor; and Section 3.2 describes the Dynamic Entrainment technique and explains how the predictor is able to keep the system in the 1:1 entrainment regime dynamically without breaking it into N:M responses.

### 3.1 Data-driven process

In this section we generate a dataset from the mechanistic model given in (1). Our focus in this part is mainly to define the predictors and responses which are included in the training dataset. In general, the deep learning model we use here, learns the relation between the input-predictors and output-responses to predict the best output-response based on the given input-predictors.

Figures 2 *A*–*C* represent the resonant response of the mesencephalic V neuron of a rat’s brainstem to pulses of injected current with a 10 ms period. In Figs. 2 *A*–*C* the forcing period increases, while the magnitude of the force remains fixed. As shown in Fig. 2*B*, if the forcing has 10 ms period, the cell is able to fire or generate an action potential. However, by decreasing or increasing the forcing period, the cell does not generate an action potential anymore as in Figs. 2 *A* and *C*. This is mainly because the forcing period does not capture the natural frequency of the cell which is known as the resonant behavior. This example illustrates that based on the first input force, the cell might not be able to spike but if the second force is applied at a right time, then the next response would be higher than the previous one and as a result, the cell membrane potential is brought up closer to its action potential threshold. By repeating this process (i.e. applying the next input force at a right time) the cell will eventually produce an action potential.

**Fig. 2:**
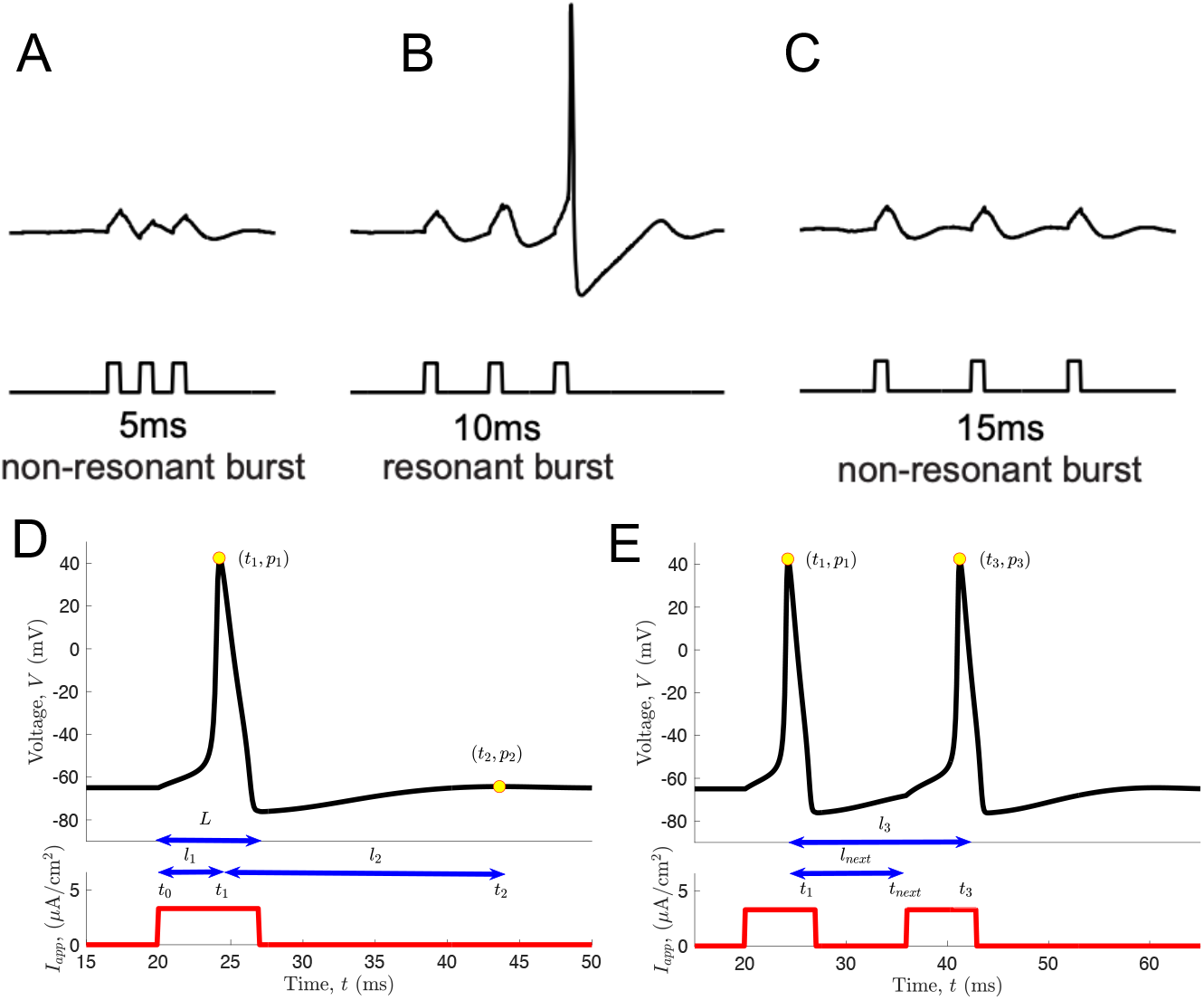
(*A*–*C*) Resonant response of the mesencephalic V neuron of a rat’s brainstem to pulses of injected current having a 10 ms period (in vitro) from [48]. From (*A*) to (*C*) the forcing period increases, while the magnitude of the force remains fixed. (*A*) The forcing period is 5 ms which is not enough to excite the cell for an action potential. (*B*) The forcing period of 10 ms brings the cell near the spiking threshold. (*C*) The forcing period is larger than the natural frequency of the cell and dies off the depolarization process. (*D* and *E*) The voltage trace *V* (*t*) versus time *t* of the mechanistic model (1) for two different injected currents *I*_*app*_. The parameter values and gating functions are *C* = 1 *µ*F/cm^2^, *g*_*Na*_ = 120 mS/cm^3^, *g*_*K*_ = 36 mS/cm^3^, *g*_*L*_ = 0.3 mS/cm^3^, *E*_*Na*_ = 55 mV, *E*_*K*_ = *−*77 mV and *E*_*L*_ = *−*54.4 mV. In both (*D*) and (*E*), *Amp* is the magnitude of the applied current or force and *L* is the active length of the force (i.e. the amount of the time the force system remains active). In (*D*), *l*_1_ = *t*_1_ *− t*_0_ is the time difference between the initiation of the force *t*_0_ and the time of the maximum voltage *t*_1_ when the force is on; *p*_1_ is the maximum voltage when the force is on; *l*_2_ = *t*_2_ *− t*_1_ is the time difference between the maximum voltage when the force is on *t*_1_ and the maximum voltage after the force turns off *t*_2_; and *p*_2_ is the maximum voltage after the force turns off. In (*E*), *l*_*next*_ = *t*_*next*_ *− t*_1_ is the time difference between the time of the maximum voltage when the first force is on *t*_1_ and the best initiation time for the second pulse to generate the second maximum voltage *t*_*next*_; *l*_3_ = *t*_3_ *− t*_1_ is the time difference between the maximum voltage for the first and second pulses *t*_1_ and *t*_3_, respectively; and *p*_3_ is the maximum voltage for the second pulse.

Predictor model attains some inputs known as input-predictors [49]. In addition, the predictor model produces output-responses which are the prediction of the model. For the data-driven processing, it is necessary to come up with an explanation of the input-predictors as well as the output-responses in a way that can be used as a synthetic dataset for the deep learning approach. By simulating the Hodgkin-Huxley model given in (1), the output is a time series data which is called voltage trace. Figures 2 *D* and *E* shows the voltage trace of (1) with the parameters given earlier in Section 2 as well as in the figure caption. Extracting more information from this time series data might help to better understand the shape of the voltage trace, however, it will increase the dimensionality of the training dataset. It should be noted that by increasing the dimensionality of the training dataset, it is necessary to have more samples which create some problems. For a dataset with higher dimension, using the same number of samples makes the dataset sparse which results in the low accuracy of the trained predictor. Another issue would be the computational costs of generating a dense training dataset, which in turn depends on how long each simulation of the mechanistic model takes to complete. The problem usually rises up with a large training dataset is that the training procedure of the predictor model can be very time consuming. In other words, more samples in the training dataset necessitates more training processes to obtain the well trained predictor.

#### 3.1.1 Input-predictors

As mentioned earlier, the role of a predictor is to estimate the output-responses based on the provided input-predictors. These input-predictors are shown in Fig. 2*D* and can be classified as the following: (1) *Amp*: the magnitude of the applied current or force; (2) *L*: the active length of the force (i.e. the amount of the time the force remains active); (3) *l*_1_ = *t*_1_ *− t*_0_: the time difference between the initiation of the force *t*_0_ and the time of the maximum voltage *t*_1_ when the force is on; (4) *p*_1_: the maximum voltage when the force is on; (5) *l*_2_ = *t*_2_ *− t*_1_: the time difference between the maximum voltage when the force is on *t*_1_ and the maximum voltage after the force turns off *t*_2_; and (6) *p*_2_: the maximum voltage after the force turns off.

#### 3.1.2 Output-responses

The ultimate goal here is to predict the best activation time of the force for the given predictors in order to have the system remains in the 1:1 entrainment regime. This can be computed numerically by activating another force with the same length and magnitude right after the first force and move it forward in time. By tracking the value of *p*_2_ which is the maximum response after the first force, we expect this value keeps increasing and after it reaches its maximum value, it will start decreasing. Therefore, we can record this turning point as the best next point to activate the force. The output-responses are shown in Fig. 2*E* and defined as: (1) *l*_*next*_ = *t*_*next*_ *− t*_1_: the time difference between the time of the maximum voltage when the first force is on *t*_1_ and the best initiation time for the second pulse to capture the second maximum voltage *t*_*next*_; (2) *l*_3_ = *t*_3_ *− t*_1_: the time difference between the maximum voltage for the first and second pulses *t*_1_ and *t*_3_, respectively; and (3) *p*_3_: the maximum voltage for the second pulse.

#### 3.1.3. Correlation matrix

Throughout the process of producing the training dataset, both the magnitude of the applied current *Amp* and the active length of the force *L* will be drawn randomly from uniform distributions. Note that for choosing the ranges of these distributions, we must consider that any values below the lower or above the upper bounds does not provide any useful information (i.e. subthreshold responses are obtained for small values of *Amp* or *L*, while spikes with a fixed peak are generated for larger values). After selecting these two input-predictors, we pass them into the mechanistic model (1) in order to obtain the rests, i.e., *l*_1_, *p*_1_, *l*_2_ and *p*_2_. This information are recorded as the input-predictors. Then, we can compute the output-responses by applying the second force pulse. Therefore, for any random values of *Amp* and *L*, the rest of the information are achieved from the mechanistic model given in (1). We repeat this process iteratively and make the comprehensive dataset to train the predictor model.

Note that before training the predictor model over a training dataset, we must check if there are high correlations between any of these defined input-predictors and/or output-responses. Figure 3 represents the correlation matrix of the input-predictors and output-responses. Here, we can observe there is a very high correlation (above 90%) between (*l*_*next*_, *l*_3_) and (*p*_1_, *p*_3_). This means preserving *l*_3_ and *p*_3_ among the output-responses would be redundant and make the process of training longer as instead of dealing with 7 dimensions we must train the predictor model over 9 dimensions. This technique helps us to reduce the dimensionality of the training dataset by keeping the minimum number of useful and necessary information. Figure 4 presents the Kernel Density Estimate (KDE) plot on the diagonal, the scatter plot on the lower triangle and 2d KDE on the upper triangle of the input-predictors *Amp, L, l*_1_, *p*_1_, *l*_2_, *p*_2_ and the output-responses *l*_*next*_, *l*_3_, *p*_3_, where *Amp* and *L* are chosen from uniform distributions with the ranges of [0.5, 15] and [5, 15], respectively. This figure describes the relationship between the input-predictors with themselves as well as the output-responses individually. As shown here, there is a high non-linearity relationship among different components of this training dataset.

**Fig. 3:**
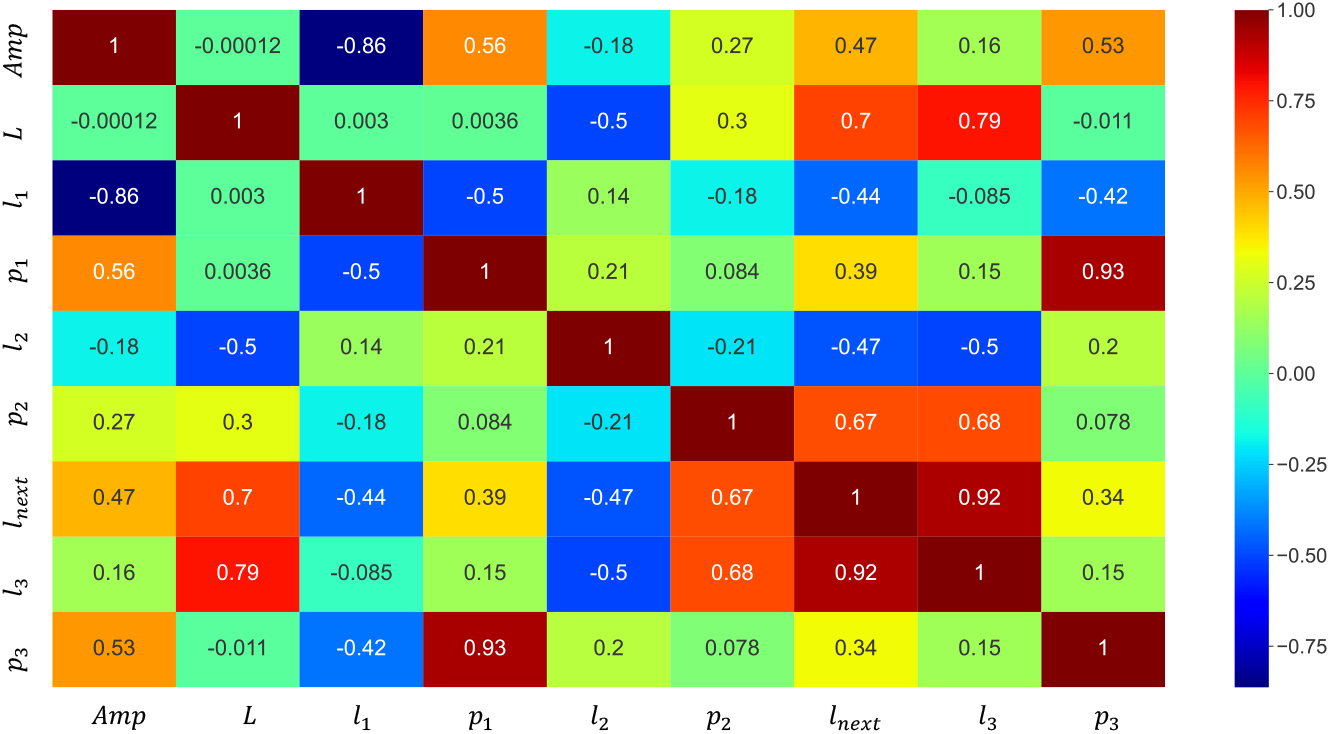
Correlation matrix of the input-predictors *Amp, L, l*_1_, *p*_1_, *l*_2_, *p*_2_ and the output-responses *l*_*next*_, *l*_3_ and *p*_3_. *Amp* and *L* are chosen from a uniform distribution with the ranges of [0.5, 15] and [5, 15], respectively.

**Fig. 4:**
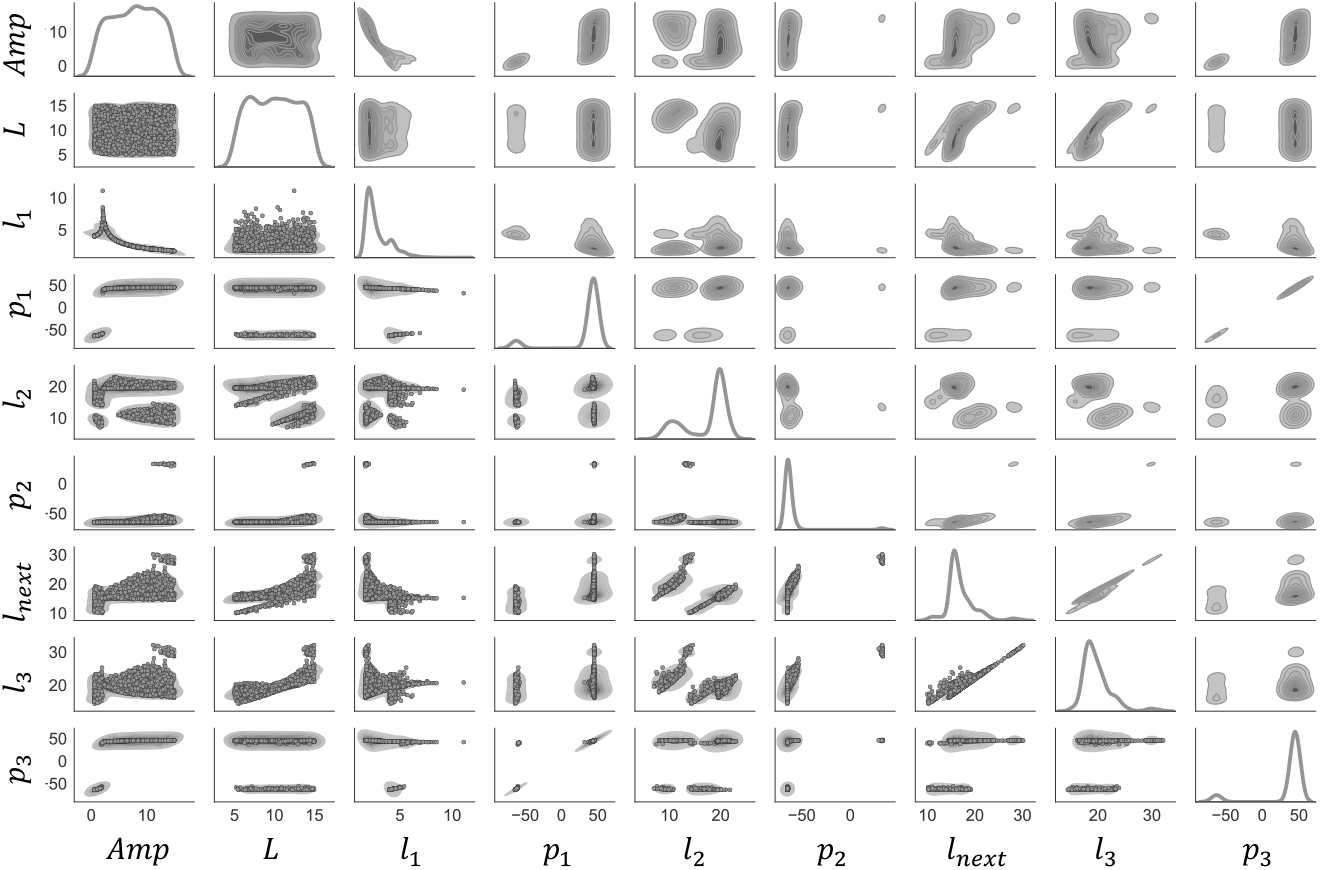
Kernel Density Estimate (KDE) plot of *∼* 200, 000 samples with input-predictors *Amp, L, l*_1_, *p*_1_, *l*_2_, *p*_2_ and the output-responses *l*_*next*_, *l*_3_, *p*_3_. *Amp* and *L* are chosen from a uniform distribution with the range of [0.5, 15] and [5, 15], respectively.

### 3.2 Dynamic entrainment

Figure 5 represents the cell membrane potential *V* (*t*) versus time *t* obtained from the mechanistic model given in (1), for several different scenarios explained below. Note that the system is kept at the steady state regime before applying the first pulse. In Figs. 5 *A*–*C*, the forcing amplitude *Amp* = 2.2 and the forcing period *T* is 18, 19 and 21, respectively. The yellow shaded regions here show that an applied pulse with amplitude *Amp* = 2.2 is strong enough to bring the cell near its firing threshold and is able to generate an action potential. In Figs. 5 *A* and *B*, there is no action potential generated by the second pulse, however an action potential occurs right after the second injected pulse in Fig. 5*C*. These observations raise a question why in Figs. 5 *A* and *B*, the second pulse with the same magnitude as the first one is not able to bring the cell above its action potential threshold anymore, while the first pulse was able to do so? To answer this question one must analyze the system behavior after depolarization state as defined and shown in Fig. 1*B*. The forcing period *T* increases in Figs. 5 *A*–*C*, therefore, the amount of time that the force system is on and off increases as well. Note that if the system has enough time after the termination of the first pulse, then the cell membrane voltage converges into its steady state regime again. At this point, the second pulse should be able to generate the next action potential. In Figs. 5 *A* and *B*, the second pulse is not applied at a right time, despite Fig. 5 *C*, in which the system remains in its 1:1 entrainment regime.

**Fig. 5:**
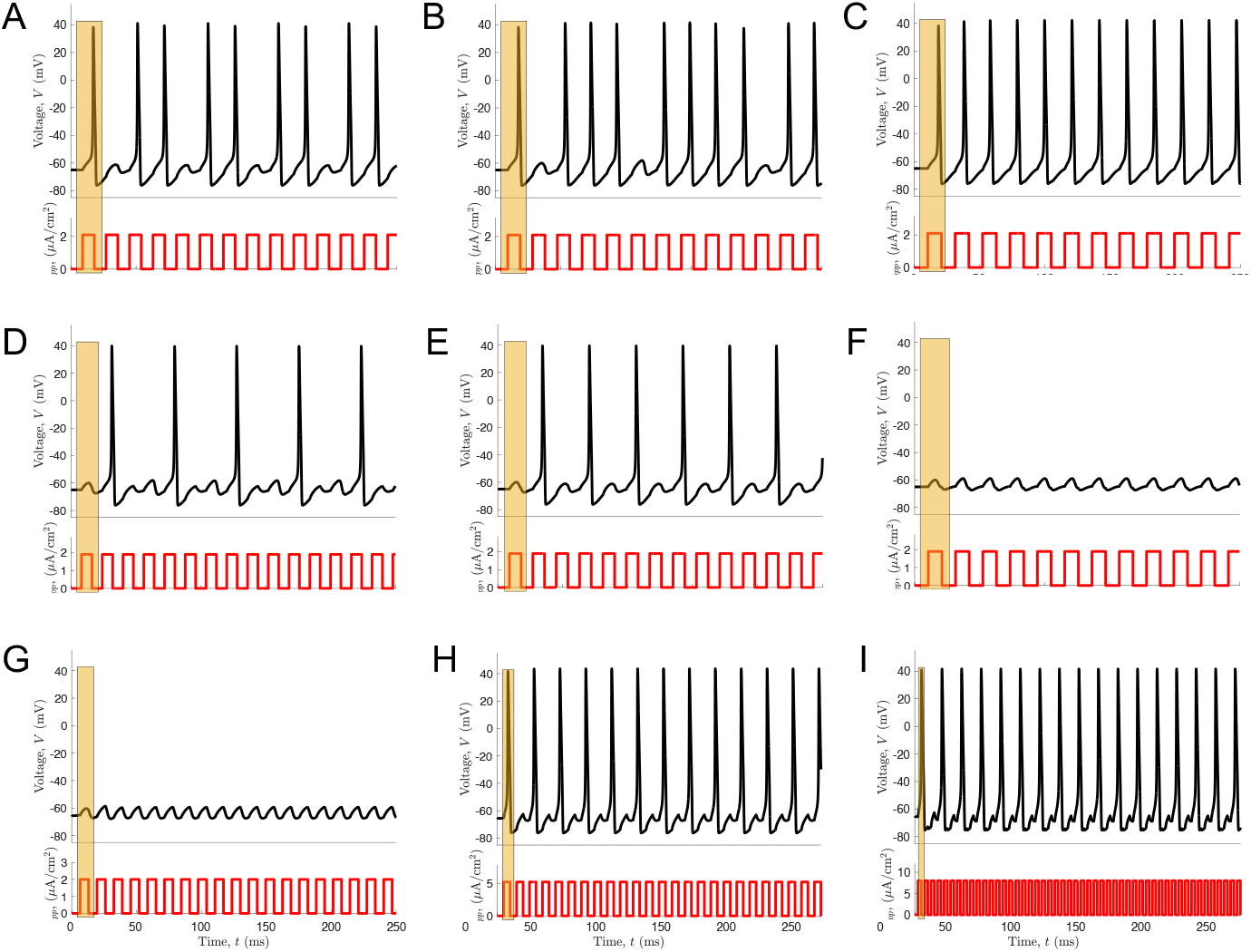
The cell membrane potential *V* (*t*) versus time *t*. The system is kept at its steady state regime before injecting the first pulse. (*A*–*C*) The forcing amplitude *Amp* = 2.2 and the forcing periods *T* = 18, 19 and 21 in (*A*), (*B*) and (*C*), respectively. The first pulse is strong enough to excite the cell to fire an action potential. In (*A*) and (*B*), the second pulse is not applied at a right time to generate an action potential, while (*C*) shows the second pulse is applied at a right time and the system remains in its 1:1 resonant entrainment regime. (*D*–*F*) The forcing amplitude is *Amp* = 1.9 and the forcing period *T* = 16, 18 and 21 in (*D*), (*E*) and (*F*), respectively. According to the yellow shaded regions, the first pulse is not strong enough to generate an action potential. In (*D*) and (*E*) the next pulses are applied at right times then action potentials occur. While (*F*) has a higher forcing period (*T* = 21), which is not aligned with the natural frequency of the cell membrane. Therefore, the system ends up with sub-threshold responses eventually. (*G*–*I*) The forcing amplitude *Amp* = 2, 5.2 and 8 and the forcing period *T* = 13, 10 and 5 in (*G*), (*H*) and (*I*), respectively. (*G*) shows sub-threshold responses, i.e. the first forcing pulse is not strong enough to generate an action potential and in addition, the forcing period is not aligned with the natural frequency of the cell, therefore the cell membrane does not generate an action potential. In (*H*) and (*I*), the first forcing pulse is strong enough to generate an action potential, however the second pulse is not being applied at a right time, therefore, we observe 2:1 and 3:1 patterns in (*H*) and (*I*), respectively.

In Figs. 5 *D*–*F*, we set *Amp* = 1.9 and the forcing period *T* is 16, 18 and 21, respectively. According to the shaded yellow regions in these figures, we can observe that the first pulse is not strong enough to generate an action potential even if the system is in its steady state regime. However, if the next pulses are applied at right times then the output voltage response will increase gradually up to a point that the cell membrane passes the firing threshold and an action potential occurs as shown in Figs. 5 *D* and *E*. On the other hand, Fig. 5 *F* has a higher forcing period (*T* = 21), which is not aligned with the natural frequency of the cell membrane, which ends up with sub-threshold responses eventually.

In Figs. 5 *A*–*F*, the effects of the forcing period *T* are studied on the cell membrane voltage trace *V* (*t*) in time *t*, while the forcing amplitude is kept constant as *Amp* = 2.2 and *Amp* = 1.9 in Figs. 5 *A*–*C* and Figs. 5 *D*–*F*, respectively. Here, we present an example to illustrate how the results might change with both the forcing period and amplitude. In Figs. 5 *G*–*I*, *Amp* = 2, 5.2, 8 and *T* = 13, 10, 5, respectively. As shown in Fig. 5*G*, we observe the cell membrane is not excited when the forcing amplitude is not strong enough. On the other hand, if the forcing amplitude increases to *Amp* = 5.2 and 8 as in Figs. 5 *H* and *I*, action potential occurs, however the forcing period *T* would be the key factor to define the voltage trace pattern. As observed, the phase is locked with 2:1 and 3:1 patterns in Figs. 5 *H* and *I* respectively, in other words, we have 2:1 and 3:1 entrainment patterns.

So far, we have provided examples in Fig. 5 to show that the strong forcing amplitude is necessary to generate action potentials but it is not sufficient to keep the system in the spiking mode. We have also shown that the forcing period dictates the voltage trace pattern. Therefore, in order to sustain a system in a 1:1 entrainment regime, the forcing pulse must be applied at a right timing window, which we call it *the window of opportunity*. Figure 6 illustrate the cell membrane potential *V* (*t*) versus time *t* obtained from the mechanistic model given in (1), with the forcing amplitude *Amp* = 2.1 and the forcing period *T* = 19. Here, the shaded yellow regions represent the window of opportunity and the blue arrows indicate the initiation of the next force. In order to remain in the 1:1 entrainment regime, the blue arrows should remain in the window of opportunity for every cycle. As shown here, since the forth pulse is not applied in the window of opportunity, no action potential occurs at that point.

**Fig. 6:**
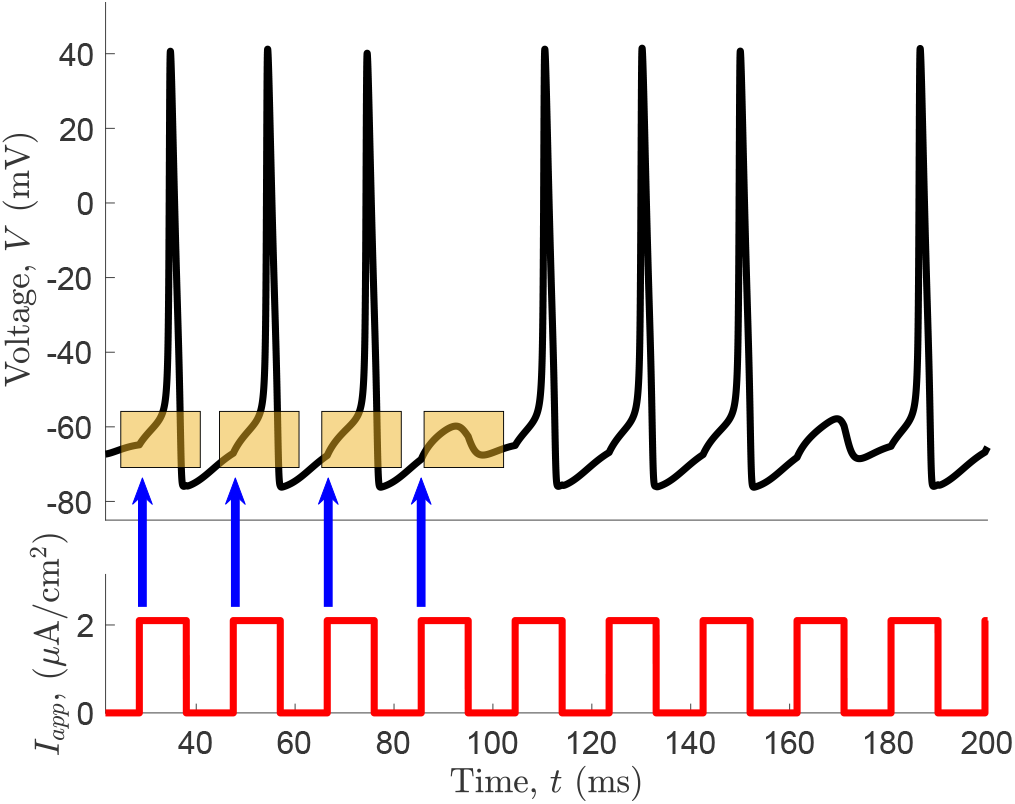
The cell membrane potential *V* (*t*) versus time *t* obtained from the mechanistic model given in (1) with the forcing amplitude *Amp* = 2.1 and the forcing period *T* = 19. Yellow shaded regions represent the window of opportunity and the blue arrows indicate the initiation of the next force. In order to remain in the 1:1 entrainment regime these blue arrows should remain in this window of opportunity for every cycle.

The length of the window of opportunity depends on the forcing amplitude *Amp*, the forcing period *T* and the length of activeness *L*. Since the voltage trace *V* (*t*) is the output of the non-linear mechanistic model given in (1), based on the input parameters *Amp* and *L*, determining the window of opportunity mathematically is very challenging. To overcome this, we use Dynamic Entrainment, which is based on data-driven process alongside the outputs of the mechanistic model for a dynamical system to train the deep learning approach in order to build a predictor to determine the best time of activeness of the force to keep the system in its 1:1 entrainment regime. Figure 7 represents the accuracy of the deep learning model prediction (*l*_*next*_) compared with the extracted feature from the output of the mechanistic model (*l*_*next*_) for a given input predictors. In other words, we compare the density estimate of the predictor model output with their true output-responses computed from the mechanistic model for the 10,000 samples drawn from the uniform distribution where the defined predictor model was not trained over. As shown in Fig. 7, the deep learning approach achieves a very good accuracy with *R*^2^ = 0.95 ^1^.

**Fig. 7:**
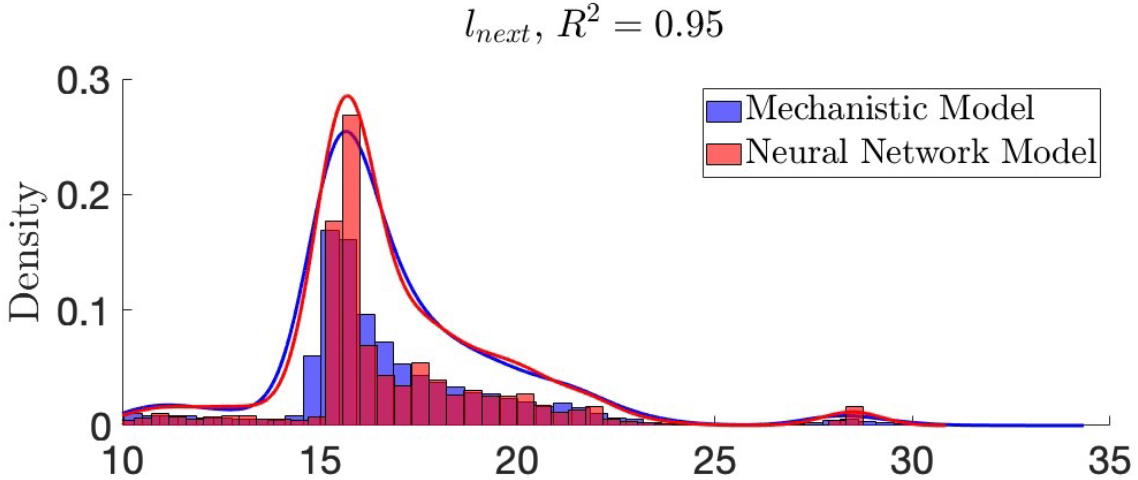
Comparison of the deep learning model and the mechanistic model (given in (1)), for the density estimate of the extracted feature *l*_*next*_ with *R*^2^ = 0.95.

Figures 8 *A, C* and *E* represent the irregular, sub-threshold and regular (i.e. many:1) cell membrane potential *V* (*t*) versus time *t*, respectively. They are obtained from the mechanistic model given in (1) with the forcing amplitude, the forcing period and the length of force activeness *Amp* = 1.9, *T* = 19 and

**Fig. 8:**
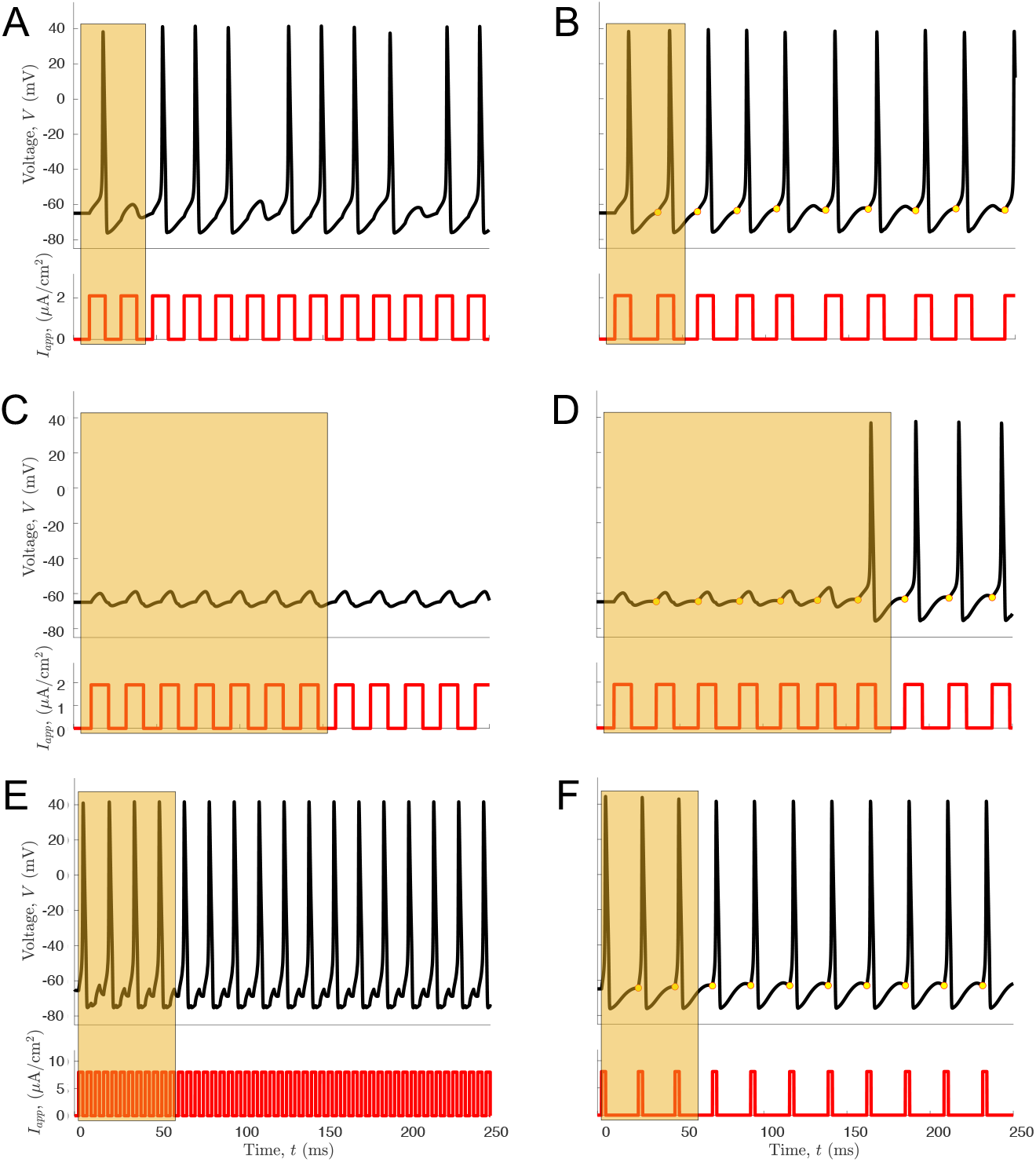
The cell membrane potential *V* (*t*) versus time *t*. (*A*), (*E*) and (*D*) obtained from the mechanistic model given in (1) with the forcing amplitude, the forcing period and the length of force activeness *Amp* = 1.9, *T* = 19 and *L* = *T/*2 = 9.5; *Amp* = 1.9, *T* = 21 and *L* = *T/*2 = 10.5; and *Amp* = 8, *T* = 5 and *L* = *T/*2 = 2.5, respectively. In (*B*), (*D*) and (*F*), the deep learning approach is applied to the information extracted from the previous voltage response to predict *l*_*next*_ to achieve 1:1 entrainment pattern for the cell membrane potential *V* (*t*) with the same forcing amplitude and length of force activeness as in (*A*), (*E*) and (*D*), respectively. Here the yellow dots represent the time (*t*_1_ + *l*_*next*_, see Fig. 2*E*) at which the next pulse must be applied in order to keep the cell membrane in the spiking mode.

*L* = *T/*2 = 9.5; *Amp* = 1.9, *T* = 21 and *L* = *T/*2 = 10.5; and *Amp* = 8, *T* = 5 and *L* = *T/*2 = 2.5, respectively. As shown in the shaded yellow region in Fig. 8 *A*, the first pulse is strong enough to generate an action potential when the system is in steady state, however a spike is missed as the second pulse is not being applied at the right time. For later times, we observe irregular pattern of missing spikes in the cell membrane potential *V* (*t*). The shaded yellow region in Fig. 8 *C* shows that the first pulse is not strong enough to generate an action potential when the system is in the steady state. In addition the forcing period is not aligned with the natural frequency of the cell membrane to generate an action potential in later times beyond the shaded yellow region. In Fig. 8 *E*, the first pulse is strong enough to generate an action potential when the system is in the steady state as shown in the shaded yellow region of this figure. However, the forcing period is not aligned with the natural frequency of the cell membrane to generate an action potential for all injected pulses. Therefore, there are missing spikes in the cell membrane potential *V* (*t*) pattern. In Figs. 8 *B, D* and *F*, the deep learning approach is applied to the information extracted from the previous voltage response to predict *l*_*next*_ to achieve 1:1 entrainment pattern for the cell membrane potential *V* (*t*) with the same forcing amplitude *Amp* and length of force activeness *L* as in Figs. 8 *A, C* and *E*, respectively. The yellow dots represent the time (*t*_1_ + *l*_*next*_, see Fig. 2*E*) at which the next pulse must be applied in order to keep the cell membrane in the spiking mode. All these yellow dots are predicted by the predictor model and they all lie in their corresponding window of opportunity. This approach is called Dynamic Entrainment as it can predict the initiation of the next force dynamically and will be able to keep the system in the 1:1 entrainment regime.

## 4 Discussion

The term “data-driven” process has become more prevalent over the past few years across nearly every fields including neuroscience [50, 51], climate science [52], disease modeling [53, 54], fluid dynamics [55, 56] and particularly in the field of dynamical systems [57–59] which focuses on discovering and characterizing dynamical systems purely from data using machine learning and deep learning approaches. The development of various algorithms in machine learning, deep learning, data science and other types of applied mathematics, statistics and optimization along with an incredible growth in the quantity of data collected from simulations and experiments, led to the emergence of the data-driven field.

In this paper, we have introduced the novel ***Dynamic Entrainment*** technique which is a type of “data-driven dynamical systems” and is combining the deep learning technique with a massive dataset extracted from a prescribed dynamical system. This technique is able to characterize the Hodgkin-Huxley model by extracting data from the voltage trace and learns to keep the system in 1:1 entrainment regime if the first input force is strong enough to generate an action potential by predicting the best initiation time for the force system. Additionally, this method is able to achieve a better reaction after each cycle in an iterative manner, resulting in an action potential in situations where the force is not strong enough to produce a spike when the system is maintained at steady state.

The Dynamic Entrainment technique opens many opportunities in many different fields. For instance, in fluid dynamics and waves, the natural frequency of ocean waves might become close to that of the movement of a ship which could be a life threatening condition for on boards. This problem can be resolved by removing the ship out of the resonant mode by applying an external force to the ship at a right time and predicting the best initiation of the force. This force can be generated by the ship engine power. The other potential future work is in cardiology where the Dynamic Entrainment technique can act as a great pace maker tool. Around 15% of all deaths in the United State are caused by Sudden Cardiac Arrest (SCA) [60], which happens when the electrical system in the heart is not working properly. In this situation, Cardiopulmonary Resuscitation (CPR) must be applied no later than a couple of minutes as the brain will be damaged due to the lack of oxygen. This procedure must be done by specialists. Without any specialist nearby, saving the patient is almost impossible as others might not be professional to be able to complete the CPR procedure properly [61]. Note that, pushing the chest too fast or too slow and not deep enough or too deep might cause the oxygen to not circulate throughout the body, which causes the death eventually. The Dynamic Entrainment technique can be helpful in this case as it might be able to assist any amature to act as efficient as a specialist to treat the SCA patients.

Last but not least, the Dynamic Entrainment technique can be applied to other neuronal models such as Morris Lecar and Fitzhugh Nagumo models. Since these models are simpler in terms of dimensionality compared to Hodgkin-Huxley model, Morris Lecar or Fitzhugh Nagumo models can help us to analyze the Dynamic Entrainment methodology in dynamical system point of view by considering the nullclines and phase plane analysis.

## Acknowledgments

S.S. acknowledges the financial support from the National Science Foundation under Grant No. DMS-2152115. P.S. gratefully acknowledges the support from the Department of Mathematics & Statistics and College of Art and Sciences from Georgia State University. S.S. and P.S. thank Drs. Igor Belykh and Casey Diekman for several helpful conversations.

## Declarations

The authors declare that they have no conflict of interest.

## Footnotes

1 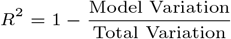

## Notes

### Competing Interest Statement

The authors have declared no competing interest.

## References

[1] Beletskii, V.: Resonance rotation of celestial bodies and cassini’s laws. Celestial Mechanics 6(3), 356–378 (1972)

[2] Landa, P.S.: Nonlinear Oscillations and Waves in Dynamical Systems vol. 360, (2013)

[3] Brown, R., Chua, L.O.: Clarifying chaos iii: Chaotic and stochastic processes, chaotic resonance, and number theory. International Journal of Bifurcation and Chaos 9(05), 785–803 (1999)

[4] Wargitsch, C., Hübler, A.: Resonances of nonlinear oscillators. Physical Review E 51(2), 1508 (1995)

[5] Andrievsky, B., Fradkov, A.: Feedback resonance in single and coupled 1-dof oscillators. International Journal of Bifurcation and Chaos 9(10), 2047–2057 (1999)

[6] Fradkov, A.L.: Feedback resonance in nonlinear oscillators. In: 1999 European Control Conference (ECC), pp. 3599–3604 (1999). IEEE

[7] Dupuit, M., de Contades, M., Ilougan, R.: Mahyer. rapport de la commission d’enquéte (*) nommée par arrété de m. le préfet de maine-et-loire, en date du 20 avril 1850, pour rechercher les causes et les circinstances qui ont amene la chute du pont suspendu de la basse-chaine. Ann. Ponts Chaussé 3130, 394–411 (1850)

[8] Belykh, I., Bocian, M., Champneys, A.R., Daley, K., Jeter, R., Macdonald, J.H., McRobie, A.: Emergence of the london millennium bridge instability without synchronisation. Nature communications 12(1), 7223 (2021)

[9] Huygens, C.: Horologium Oscillatorium, (1980)

[10] Oliveira, H.M., Melo, L.V.: Huygens synchronization of two clocks. Scientific reports 5(1), 1–12 (2015)

[11] Blekhman, I.I.: Synchronization in Science and Technology, (1988)

[12] Cahn, C.: Review of’synchronization systems in communication and control’(lindsey, wc; 1972). IEEE Transactions on Information Theory 19(5), 714–715 (1973)

[13] Fradkov, A.L., Andrievsky, B.: Synchronization and phase relations in the motion of two-pendulum system. International Journal of Non-Linear Mechanics 42(6), 895–901 (2007)

[14] Tyrrell, A., Auer, G., Bettstetter, C.: Firefly synchronization in ad hoc networks. In: Proceedings of the MiNEMA Workshop (2006)

[15] Moshal, K.S., Camel, C.K., Kartha, G.K., Steed, M.M., Tyagi, N., Sen, U., Kang, Y.J., Lominadze, D., Maldonado, C., Tyagi, S.C.: Cardiac dys-synchronization and arrhythmia in hyperhomocysteinemia. Current neurovascular research 4(4), 289–294 (2007)

[16] Mase, M., Faes, L., Antolini, R., Scaglione, M., Ravelli, F.: Quantification of synchronization during atrial fibrillation by shannon entropy: validation in patients and computer model of atrial arrhythmias. Physiological measurement 26(6), 911 (2005)

[17] Casula, E.P., Mayer, I.M., Desikan, M., Tabrizi, S.J., Rothwell, J.C., Orth, M.: Motor cortex synchronization influences the rhythm of motor performance in premanifest huntington’s disease. Movement Disorders 33(3), 440–448 (2018)

[18] Wu, H.-M., Hsiao, F.-J., Chen, R.-S., Shan, D.-E., Hsu, W.-Y., Chiang, M.-C., Lin, Y.-Y.: Attenuated nogo-related beta desynchronisation and synchronisation in parkinson’s disease revealed by magnetoencephalo-graphic recording. Scientific reports 9(1), 1–12 (2019)

[19] Arbel-Goren, R., Buonfiglio, V., Di Patti, F., Camargo, S., Zhitnitsky, A., Valladares, A., Flores, E., Herrero, A., Fanelli, D., Stavans, J.: Robust, coherent, and synchronized circadian clock-controlled oscillations along anabaena filaments. Elife 10, 64348 (2021)

[20] Beer, K., Joschinski, J., Arrazola Sastre, A., Krauss, J., Helfrich-Förster, C.: A damping circadian clock drives weak oscillations in metabolism and locomotor activity of aphids (acyrthosiphon pisum). Scientific reports 7(1), 14906 (2017)

[21] Liao, G., Diekman, C., Bose, A.: Entrainment dynamics of forced hierarchical circadian systems revealed by 2-dimensional maps. SIAM Journal on Applied Dynamical Systems 19(3), 2135–2161 (2020)

[22] Creaser, J.L., Diekman, C.O., Wedgwood, K.C.: Entrainment dynamics organised by global manifolds in a circadian pacemaker model. Frontiers in Applied Mathematics and Statistics, 52 (2021)

[23] Diekman, C.O., Bose, A.: Reentrainment of the circadian pacemaker during jet lag: East-west asymmetry and the effects of north-south travel. Journal of theoretical biology 437, 261–285 (2018)

[24] Wulff, K., Gatti, S., Wettstein, J.G., Foster, R.G.: Sleep and circadian rhythm disruption in psychiatric and neurodegenerative disease. Nature Reviews Neuroscience 11(8), 589–599 (2010)

[25] Guevara, M.R., Glass, L.: Phase locking, period doubling bifurcations and chaos in a mathematical model of a periodically driven oscillator: A theory for the entrainment of biological oscillators and the generation of cardiac dysrhythmias. Journal of mathematical biology 14(1), 1–23 (1982)

[26] Hayashi, C., Shibayama, H., Nishikawa, Y.: Frequency entrainment in a self-oscillatory system with ex-ternal force. IRE Transactions on Circuit Theory 7(4), 413–422 (1960)

[27] Hirschie Johnson, C., Elliott, J.A., Foster, R.: Entrainment of circadian programs. Chronobiology international 20(5), 741–774 (2003)

[28] Kopell, N., Ermentrout, G.B.: Mechanisms of phase-locking and frequency control in pairs of coupled neural oscillators. Handbook of dynamical systems 2, 3–54 (2002)

[29] Medvedev, A., Mattsson, P., Zhusubaliyev, Z.T., Avrutin, V.: Nonlinear dynamics and entrainment in a continuously forced pulse-modulated model of testosterone regulation. Nonlinear dynamics 94(2), 1165–1181 (2018)

[30] Pikovsky, A., Rosenblum, M., Kurths, J.: Synchronization: a universal concept in nonlinear science. American Association of Physics Teachers (2002)

[31] Pittendrigh, C.S., Minis, D.H.: The entrainment of circadian oscillations by light and their role as photoperiodic clocks. The American Naturalist 98(902), 261–294 (1964)

[32] Roenneberg, T., Daan, S., Merrow, M.: The art of entrainment. Journal of biological rhythms 18(3), 183–194 (2003)

[33] Winfree, A.T.: Biological rhythms and the behavior of populations of coupled oscillators. Journal of theoretical biology 16(1), 15–42 (1967)

[34] Thaut, M.H., McIntosh, G.C., Hoemberg, V.: Neurobiological foundations of neurologic music therapy: rhythmic entrainment and the motor system. Frontiers in psychology 5, 1185 (2015)

[35] Khan, E., Saghafi, S., Diekman, C.O., Rotstein, H.G.: The emergence of polyglot entrainment responses to periodic inputs in vicinities of hopf bifurcations in slow-fast systems. Chaos: An Interdisciplinary Journal of Nonlinear Science 32(6), 063137 (2022)

[36] Hodgkin, A.L., Huxley, A.F.: A quantitative description of membrane current and its application to conduction and excitation in nerve. The Journal of physiology 117(4), 500–544 (1952)

[37] Nikookar, S., Sakharkar, P., Smagh, B., Amer-Yahia, S., Roy, S.B.: Guided task planning under complex constraints. In: 2022 IEEE 38th International Conference on Data Engineering (ICDE), pp. 833–845 (2022). IEEE

[38] Kober, J., Bagnell, J.A., Peters, J.: Reinforcement learning in robotics: A survey. The International Journal of Robotics Research 32(11), 1238–1274 (2013)

[39] Daoun, D., Ibnat, F., Alom, Z., Aung, Z., Azim, M.A.: Reinforcement learning: a friendly introduction. In: The International Conference on Deep Learning, Big Data and Blockchain (Deep-BDB 2021), pp. 134–146 (2022). Springer

[40] Nikookar, S., Sakharkar, P., Somasunder, S., Basu Roy, S., Bienkowski, A., Macesker, M., Pattipati, K.R., Sidoti, D.: Cooperative route planning framework for multiple distributed assets in maritime applications. SIGMOD 2022 (2022)

[41] Gamboa, J.C.B.: Deep learning for time-series analysis. arXiv preprint 1701.01887 (2017)

[42] Busseti, E., Osband, I., Wong, S.: Deep learning for time series modeling. Technical report, Stanford University, 1–5 (2012)

[43] Fu, Y., Wu, D., Boulet, B.: Reinforcement learning based dynamic model combination for time series forecasting. In: Proceedings of the AAAI Conference on Artificial Intelligence, vol. 36, pp. 6639–6647 (2022)

[44] Wu, T., Ortiz, J.: Rlad: Time series anomaly detection through reinforcement learning and active learning. arXiv preprint 2104.00543 (2021)

[45] Kutz, J.N.: Data-driven Modeling & Scientific Computation: Methods for Complex Systems & Big Data, (2013)

[46] Bielza, C., Larraũaga, P.: Data-driven Computational Neuroscience: Machine Learning and Statistical Models, (2020)

[47] Ermentrout, G.B., Terman, D.H.: Mathematical Foundations of Neuroscience vol. 35, (2010)

[48] Izhikevich, E.M.: Dynamical Systems in Neuroscience. MIT press, ??? (2007)

[49] Brunton, S.L., Kutz, J.N.: Data-driven Science and Engineering: Machine Learning, Dynamical Systems, and Control, (2022)

[50] Van Horn, J.D.: Bridging the brain and data sciences. Big Data 9(3), 153–187 (2021)

[51] Stougiannis, A., Pavlovic, M., Tauheed, F., Heinis, T., Ailamaki, A.: Data-driven neuroscience: enabling breakthroughs via innovative data management. In: Proceedings of the 2013 ACM SIGMOD International Conference on Management of Data, pp. 953–956 (2013)

[52] Krčál, L., Ho, S.-S.: A scidb-based framework for efficient satellite data storage and query based on dynamic atmospheric event trajectory. In: Proceedings of the 4th International ACM SIGSPATIAL Workshop on Analytics for Big Geospatial Data, pp. 7–14 (2015)

[53] Adiga, A., Chen, J., Marathe, M., Mortveit, H., Venkatramanan, S., Vullikanti, A.: Data-driven modeling for different stages of pandemic response. Journal of the Indian Institute of Science 100(4), 901–915 (2020)

[54] Al-Mamun, M.A., Smith, R.L., Nigsch, A., Schukken, Y.H., Gröhn, Y.T.: A data-driven individual-based model of infectious disease in livestock operation: A validation study for paratuberculosis. PLoS One 13(12), 0203177 (2018)

[55] Xie, J., Mao, S., Zhang, Z., Liu, C.: Data-driven approaches for characterization of aerodynamics on super high-speed elevators. Journal of Computing and Information Science in Engineering 23(3), 031004 (2023)

[56] Gebraad, P.M., Teeuwisse, F.W., van Wingerden, J.-W., Fleming, P.A., Ruben, S.D., Marden, J.R., Pao, L.Y.: A data-driven model for wind plant power optimization by yaw control. In: 2014 American Control Conference, pp. 3128–3134 (2014). IEEE

[57] Wan, Z.Y., Vlachas, P., Koumoutsakos, P., Sapsis, T.: Data-assisted reduced-order modeling of extreme events in complex dynamical systems. PloS one 13(5), 0197704 (2018)

[58] Klus, S., Nüske, F., Koltai, P., Wu, H., Kevrekidis, I., Schütte, C., Noé, F.: Data-driven model reduction and transfer operator approximation. Journal of Nonlinear Science 28, 985–1010 (2018)

[59] Okuno, S., Ikeuchi, K., Aihara, K.: Practical data-driven flood forecasting based on dynamical systems theory. Water Resources Research 57(3), 2020–028427 (2021)

[60] Adabag, A.S., Luepker, R.V., Roger, V.L., Gersh, B.J.: Sudden cardiac death: epidemiology and risk factors. Nature Reviews Cardiology 7(4), 216–225 (2010)

[61] Idris, A.H., Becker, L.B., Ornato, J.P., Hedges, J.R., Bircher, N.G., Chandra, N.C., Cummins, R.O., Dick, W., Ebmeyer, U., Halperin, H.R., et al.: Utstein-style guidelines for uniform reporting of laboratory cpr research: a statement for healthcare professionals from a task force of the American heart association, the american college of emergency physicians, the american college of cardiology, the european resuscitation council, the heart and stroke foundation of canada, the institute of critical care medicine, the safar center for resuscitation research, and the society for academic emergency medicine. Circulation 94(9), 2324–2336 (1996)

